# Extensive genomic diversity among *Mycobacterium marinum* strains revealed by whole genome sequencing

**DOI:** 10.1101/249532

**Authors:** Sarbashis Das, B. M. Fredrik Pettersson, Phani Rama Krishna Behra, Amrita Mallick, Martin Cheramie, Lisa Shirreff, Tanner DuCote, Santanu Dasgupta, Don G. Ennis, Leif. A. Kirsebom

## Abstract

*Mycobacterium marinum* is the causative agent for the tuberculosis-like disease mycobacteriosis in fish and skin lesions in humans. Ubiquitous in its geographical distribution, *M. marinum* is known to occupy diverse fish as hosts. However, information about its genomic diversity is limited. Here, we provide the genome sequences for 15 *M. marinum* strains isolated from infected humans and fish. Comparative genomic analysis of these and four available genomes of the *M. marinum* strains M, E11, MB2 and Europe reveal high genomic diversity among the strains, leading to the conclusion that *M. marinum* should be divided into two different clusters, the “M”- and the “Aronson”-type. We suggest that these two clusters should be considered, if not two separate species, at least two *M. marinum* subspecies. Our data also show that the *M. marinum* pan-genome for both groups is open and expanding and we provide data showing high number of mutational hotspots in *M. marinum* relative to other mycobacteria such as *Mycobacterium tuberculosis*. This high genomic diversity might be related to that *M. marinum* occupy different ecological niches.

## Background

The genus *Mycobacterium* comprises more than 177 species including, pathogenic, opportunistic pathogens and non-pathogenic environmental species. Mycobacteria are free-living, acid fast, robust organisms, which can sustain themselves, and even thrive, in widely diverse environments ranging from soil and tap water to animals and humans. Of interest to this study, *Mycobacterium marinum* (*Mma*) was first described in 1926 by Joseph D. Aronson (Aronson 1926). The bacterium was isolated from infected fish suffering from mycobacteriosis exhibiting similarities to tuberculosis in humans. Later, it was shown that *Mma* could also infect humans and cause skin lesions at body extremities (Sette et al. 2015; Johnson and Stout 2015). Phylogenetic analysis based on the 16S rRNA gene sequences suggested that *Mycobacterium tuberculosis* (*Mtb*) and *Mycobacterium ulcerans* are its closest neighbours (Stinear et al. 2000) and *Mma* is a member of the *M. ulcerans* clade. Together with *M. ulcerans* it constitutes the *M. ulcerans*-*M. marinum* complex, MuMC (Aronson 1926; Qi et al. 2009; Pidot et al. 2010; Kurokawa et al. 2012).

The complete genome of the *M. marinum* M strain was released 2008. The genome size is 6.5 Mb, which is 2.1 Mb longer than the *Mtb* genome. Comparison studies showed that *Mma* and *Mtb* have more than 85% genome similarity and share major virulence factors (Stinear et al. 2008). The *M. ulcerans* genome is roughly one Mb smaller than the *Mma* M strain genome (Käser et al. 2007) but their genomes are highly similar (Doig et al. 2012). They share a common ancestor and available genetic data suggest that *M. ulcerans* diverged from *M. marinum* (Röltgen et al. 2012). The MuMC and the evolution of *M.ulcerans* and its capacity to produce the toxin mycolactone have been studied using comparative genomic approaches (Käser et al. 2007; Doig et al. 2012). However, only a few *Mma* genomes are available and therefore knowledge of the genomic diversity as well as the global *Mma* gene repository is rather limited. Hence, to understand the evolutionary relationship and genomic diversity of *Mma* we decided to determine the genome sequences of strains isolated from different sources. In our analysis we also included four published *Mma* genomes of the Europe (acc no. ANPL00000000), MB2 (acc no. ANPM01000000), E11 (acc no. HG917972.2; Ummels et al. 2014), and M strains (GCA_000018345; Stinear et al. 2008).

Here we provide genomic data for 15 different *Mma* strains including type strains and isolates from infected fishes and humans from different geographical regions. Comparative analysis of 19 genomes suggests two distinct *Mma* types (or lineages) that share a common ancestor. This raises the question that the lineage to which the *Mma* M strain belongs and the other lineage, referred to as the “Aronson-lineage”, should be considered as two separate species (or subspecies). Consistent with this notion, during the course of evolution, “Aronson-lineage” members acquired an additional ribosomal operon likely through duplication. We have identified the presence of plasmid sequences, various IS elements and their distribution in the different strains, as well as sequences of phage origin and their translocation in the different strains. Additionally, we characterized the *Mma* pan-genome, the phylogenetic relationship and identified mutational hot spots. Altogether, these data provide insight into the evolutionary mechanisms of mycobacterial strain diversification.

## Results

Whole genome sequencing was performed for 15 *Mma* strains (Table 1). Of these, the type strains *Mma* CCUG20998 (hereafter referred to as CCUG) and *Mma* 1218R (referred to as 1218R) were sequenced using Pacific Biosciences (PacBio) technology and the remaining 13 strains with Illumina sequencing technology. In addition, in our comparative analysis we included four published *Mma* genome sequences referred to as M, E11, Europe and MB2 (Ummels et al. 2014). The genomes of the M and E11 strains are complete while the other two are draft genomes.

**Table 1.**
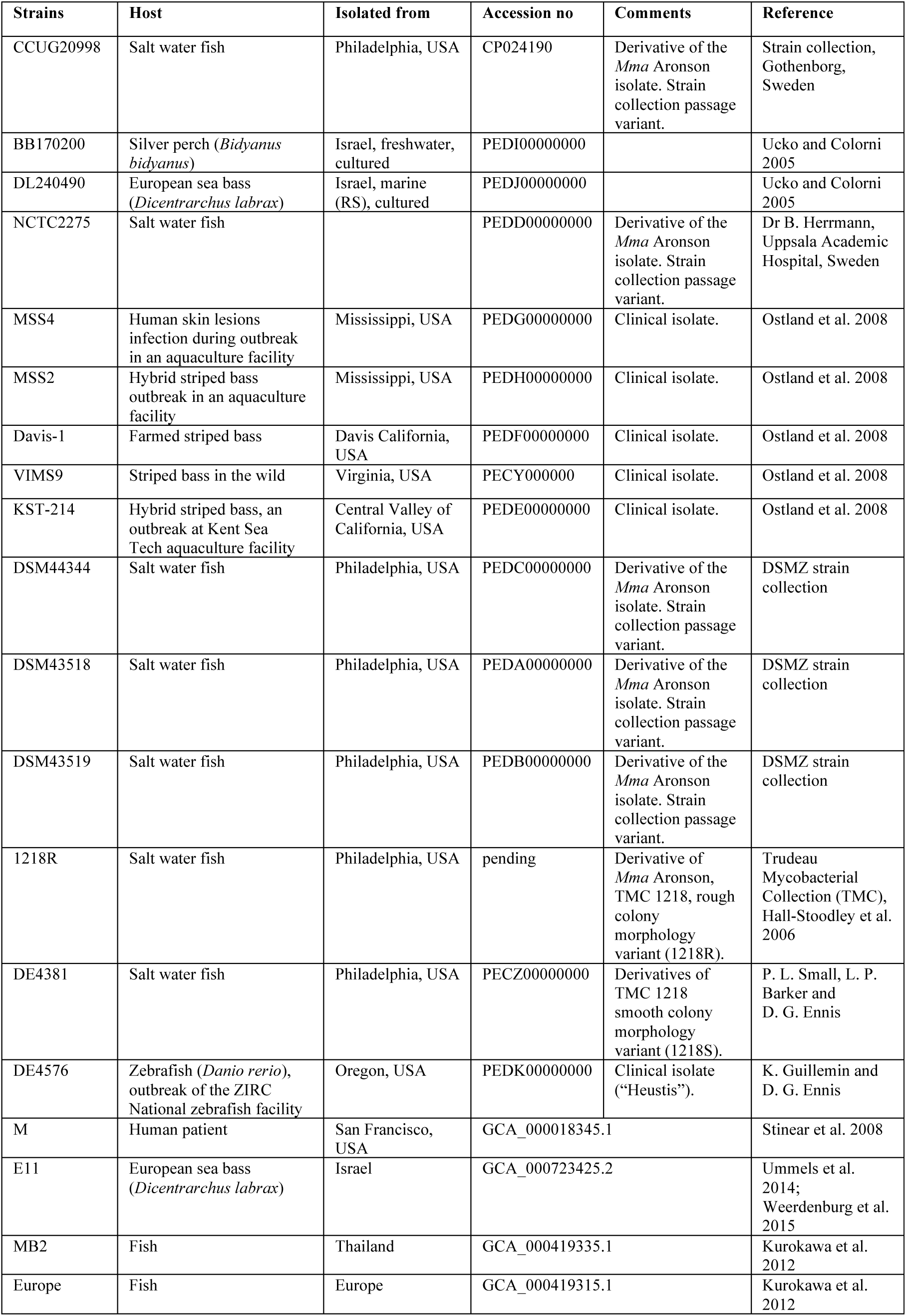
List of the *Mma* strains and corresponding sources of isolation.

## Overview of genomic features

*De novo* assembly of the long reads (average length 10 kb) derived from PacBio platform generated complete genomes (CCUG and 1218R) comprised of single scaffolds while assembly of the Illumina sequencing reads (average length 100 bp) for the other *Mma* strains resulted in near complete genomes split into multiple scaffolds. Sequencing read statistics, average read depths and assembly qualities are shown in Fig S1. The average GC-content is 65%, in keeping with the 65.7% determined for the M strain (Stinear et al. 2008) while the genome length ranged from 5.7 Mb (DL240490) to 6.6 Mb (M) representing 5343 to 5573 coding sequences (CDS; Fig 1a and Table S1). The number of predicted tRNA genes varied between 46 and 53 with 1218R having the highest number. Interestingly, the complete genomes of three strains (1218R, CCUG and E11) carry six ribosomal genes, corresponding to two ribosomal RNA operons (rRNA; 16S rRNA, 23S rRNA and 5S rRNA), while the M strain carry only one rRNA operon (Fig 1b). For the draft genomes, we predicted the presence of two 5S rRNA genes for the strains closely related to CCUG and 1218R (see below) while for the others only one was predicted (see discussion). No tRNA genes were predicted within any of the rRNA operons. Depending on strain, we also predicted the presence of 45 to 74 non-coding (nc) RNA genes (Table S1).

**Figure 1.**
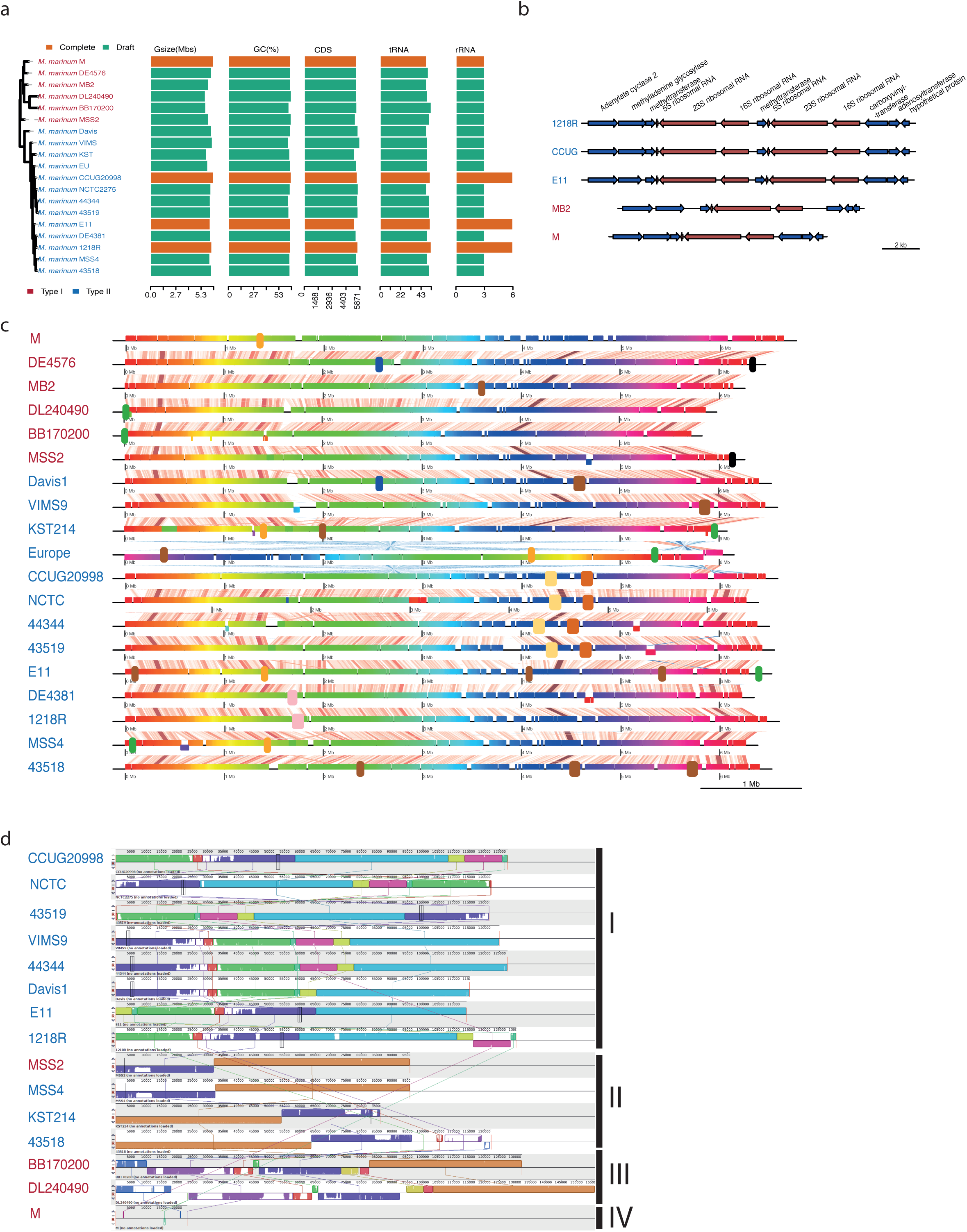
Overview of genomic features and genome alignments in *Mma* strains. (a) Bar plot showing genome size, GC-content (%), number of CDS, number of tRNA and number of rRNA in different strains along with the phylogenetic relationship. In the phylogenetic tree the strain names are in two colours representing the cluster-I (red) and cluster-II (blue) members. Complete and draft genomes are coloured by orange and green, respectively. (b) Synteny for the rRNA genes, *rrnA* and *rrnB*, present in cluster-I and cluster-II strains. Arrows represent genes and strand information. Right and left arrows indicate positive and negative strands. Red arrows refer to the rRNA operons and blue arrows mark flanking genes. (c) Whole-genome alignment of the 19 *Mma* strains where each of the coloured horizontal blocks represents one genome and the vertical bars represent homologous regions. Diagonal lines represent genomic rearrangements, whereas white gaps represent insertions/deletions. Presence of phage sequences are marked as large blocks in blue, green, yellow, and black. The same colour (except the black blocks) indicates that the phage fragments are the same while black blocks mark non-conserved phage sequences. (d) Alignment of the plasmid scaffolds in different *Mma* strain. Homologous regions in the plasmids are indicated by same coloured blocks connected with vertically lines. Partially filled regions and white regions in the blocks represent less similar sequence or unique regions respectively. All the plasmids are classified into four classes as indicated on the right side, see also the main text.

Whole genome alignment for the 19 genomes suggested that the genomes are well conserved without major genomic rearrangements (Fig 1c). However, genomic variations were apparent in two genomic regions, the “1.5-2 Mb” and “3.5-4.5 Mb” regions, in all the strains. A likely reason for this variation is the presence of different phage sequences in these regions. For example, the M strain carries a fragment in the 1.4 Mb region, which is conserved in E11, MSS4, and KST but absent in the other strains. Similarly, CCUG, NCTC2275, DSM44344 and DSM43519 have two conserved prophages located at 4.3 Mb and 4.7 Mb. The DSM43518 strain was predicted to have three prophages. Two of these (located at 4.5 Mb and 5.5 Mb) were also detected in two other strains, the “4.5 Mb-phage” in Davis-1 and the “5.5 Mb-phage” in VIMS-9 (Fig 1c). Moreover, two inversions were detected in the Europe genome and one of these covers nearly 5.5 Mb. Genome alignment, however, suggests that this inversion is likely the result of scaffolding of the contigs.

With the exception of DE4381 and DE4576, the 15 new *Mma* genomes were predicted to have plasmid sequences (Table S2; see Methods). Additionally, no plasmid was present in the MB2 and Europe strains, while CCUG and 1218R harboured complete circular plasmids encompassing 127 kb and 130 kb, respectively. For the draft genomes, DSM43519 was predicted to have the largest plasmid of 181 kb encoding 161 genes. The average GC-content for the plasmid scaffolds is 63.9%, which is lower than for the chromosomal sequences (see above) and the plasmid present in the M strain (67.9%). Plasmid alignment revealed that plasmids/ plasmid sequences could be grouped into four types: (I) CCUG, NCTC2275, DSM43519, VIMS-9, DSM44344, Davis-1 and 1218R carry the same plasmid, which is similar to the plasmid present in E11, (II) present in MSS2 and MSS4, (III) BB170200 and DL240490 have similar plasmid fragments and the sequences show high similarity compared to the pMUM003 plasmid previously reported to be present *Mma* DL240490 (Pidot et al. 2008; see discussion), and (IV) pMM23 is present only in the M strain (Fig 1d; Stinear et al. 2008). Interestingly, the type (I) plasmid carries genes encoding for two secretion systems, type IV and type VII, where the type VII genes show high homology to the ESX-5 category. The significance of the presence of these genes remains to be studied but noteworthy, ESX-5 has been suggested to be associated with slow growing pathogenic mycobacteria and to have an impact on virulence (Gröschel et al. 2016; Bosserman and Champion 2017).

## Average nucleotide identity revealed two distinct *M. marinum* strain clusters

The average nucleotide identity (ANI) value, which is useful for discriminating between species and strains (Richter and Rosselló-Móra 2009), was calculated pairwise for the homologous regions in the 19 *Mma* genomes and the two phylogenetically closest neighbours *M. ulcerans* and *Mycobacterium liflandii*. Although the ANI values for any pairs are higher than 97% (Fig 2a), hierarchical clustering on the basis of the ANI values resulted in two clusters: cluster I including the M, DE4576, MB2, DL240490, BB170200 and MSS2 strains while cluster II encompasses the remaining strains including E11, 1218R and CCUG (Fig 2a and b). Strains belonging to cluster II show significant similarity (>98.5% ANI score), while for cluster I strains the ANI scores range from 97 to 99%. We interpret this difference to reflect higher genomic diversity among cluster I members. Including *M. ulcerans* and *M. liflandii* revealed high ANI scores comparing either of these two species with several of the strains in cluster I, *e.g.,* ≈99% comparing *M. liflandii* and BB170200 or DL240490. Hence, it appears that *M. ulcerans* and *M. liflandii* are evolutionarily closer to cluster I strains than to cluster II members. In summary, *Mma* strains can be grouped into two clusters, albeit that all the ANI scores are high and above species threshold (97%; Richter and Rosselló-Móra 2009).

**Figure 2.**
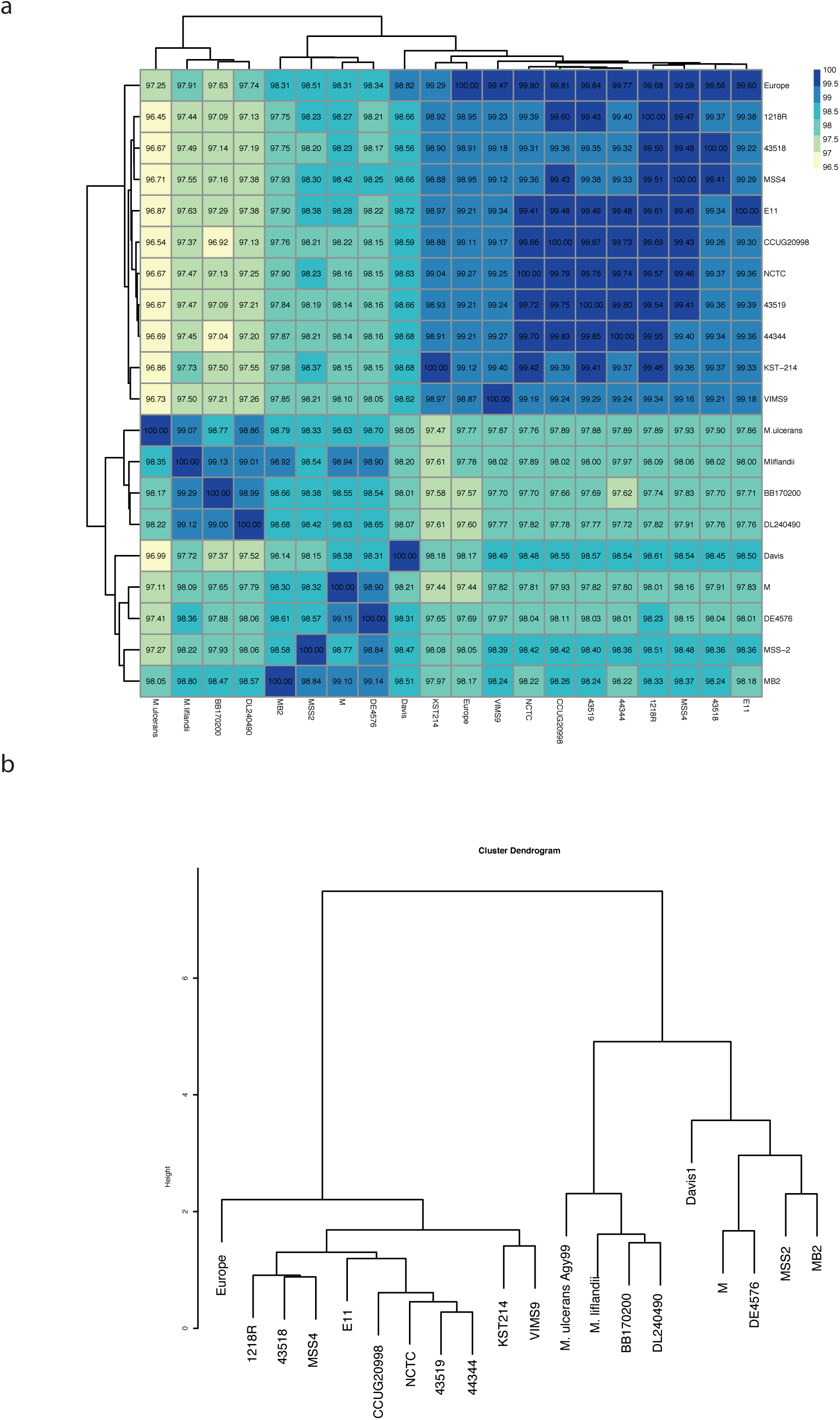
Clustering of *Mma* strains based on Average Nucleotide Identity (ANI) (a) Heat map showing ANI values for all versus all *Mma* strains including M. ulcerans and *M. liflandii.* (b) ANI values were clustered using unsupervised hierarchical clustering and plotted as dendogram.

## Cluster I and II pan-genome and core-genome sizes are different

The pan-genome includes all genes identified in all members of a species while the core-genome represents the set of genes present in all species members (Lapierre and Gogarten 2009). We used power law regression analysis and

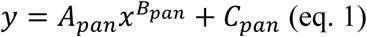

to model the distribution of the pan-genome where y = pan-genome size, x = number of genome, *A_pan_*, *B_pan_* and *C_pan_* are fitting parameters. Similarly, the core genome was modelled using

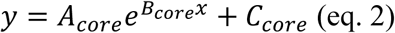

where *A_core_*, *B_core_* and *C_core_* are fitting parameters and x = number of genome. The data are shown in Fig 3a.

**Figure 3.**
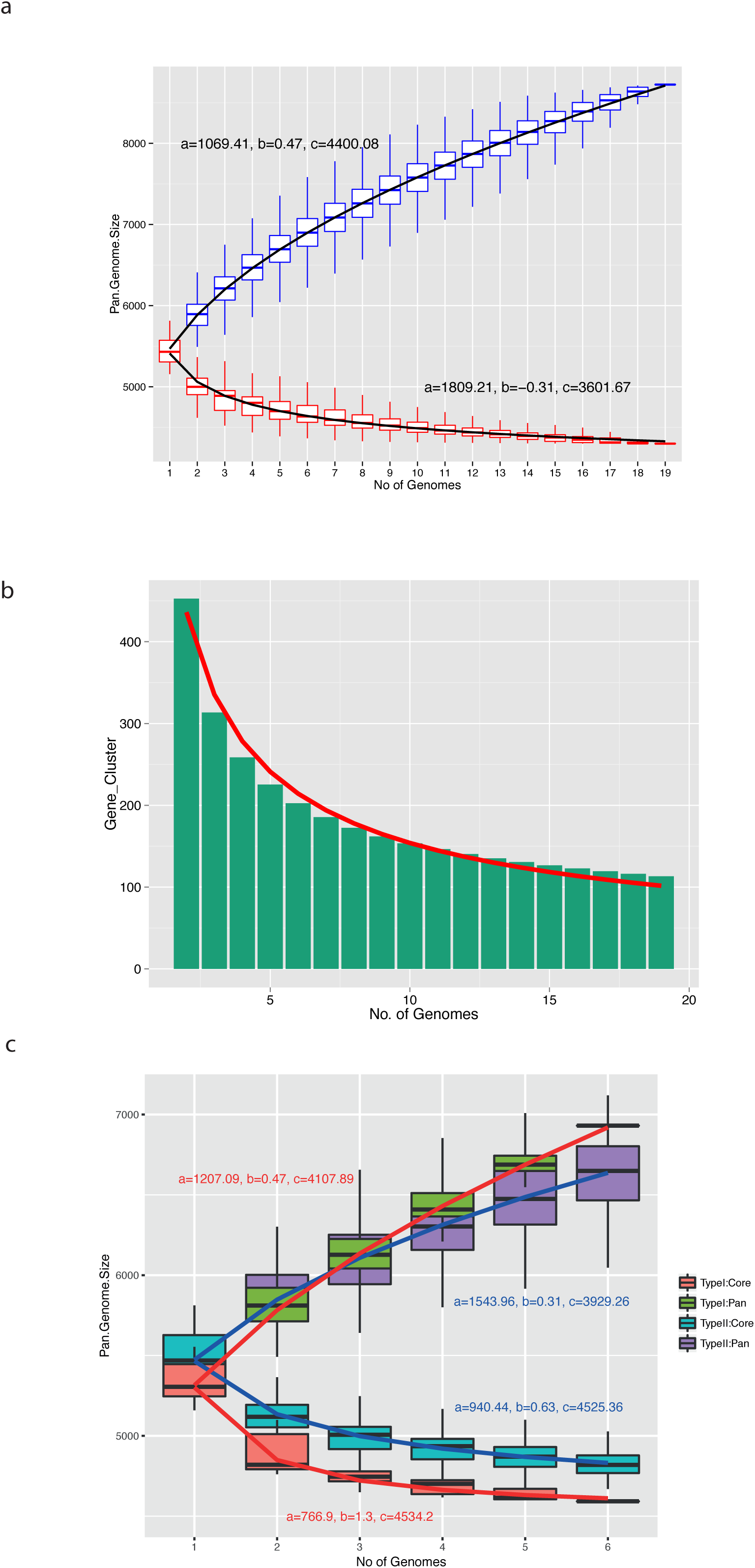
Pan-genome and core-genome of *Mma*. (a) Boxplots showing pan-genome (blue) and core-genome (red) for progressively increasing number of genomes. The black line is a fitted-line model by a regression formula (see text for details). (b) The number of new genes identified with increasing number of genomes. The red line is a fitted-line model generated by regression analysis (see text for details). (c) Similar plot as in (a) showing the results for cluster-I and cluster-II genomes separately.

For the pan-genome, *B_pan_* should be close to 1 if it is closed, which means that sequencing of additional genomes would not add any new gene to the gene repository. However, the *B_pan_* value is 0.47 suggesting that the *Mma* pan-genome is open and evolving. Moreover, based on the 19 *Mma* genomes the pan-genome size is 8725, while the core-genome comprises 4300 genes. From Fig 3b, we also estimated that approximately 100 “new” genes, not present in any of the current strains, would be identified upon sequencing an additional *Mma* genome.

Calculation of the pan- and core-genomes for cluster-I and cluster-II members revealed that cluster-I strains share 85% of their genes while any six cluster-II members share 89% (for a reliable comparison we analysed any six cluster-II strains since only six strains belong to cluster-I). Moreover, pan- and core-genome curves for cluster-I are slightly more separated than the corresponding curves for genomes belonging to cluster-II, suggesting modestly higher genomic variation among cluster-I strains compared to cluster-II members (Fig 3c; see also above).

## Variation of IS elements in *M. marinum*

Insertion (IS) elements are important factors responsible for genomic variations and dynamics (Siguier et al. 2014). The M strain carries multiple copies of eight different IS element types referred to as ‘ISMyma1-7,11’ (Stinear et al. 2008), color-coded as shown in Fig 4a. Hence, we first identified the IS elements present in the four complete genomes using ISsaga (IS-semi automated annotation; Varani et al. 2011). Subsequently, we predicted the copy number and distribution of these IS elements in the draft genomes using raw reads (see Methods); the number of IS elements of each type is indicated by lengths and coloured patches (Fig 4a). The MSS4 strain carried the highest number of predicted IS elements (n = 44) encompassing six “IS-types” (excluding ISMyma4 and ISMyma11). Of these, 11 copies were classified as ISMyma2 and 18 as ISMyma7. Moreover, the ISMyma4 type was only detected in the M, DE4576 and Davis1 strains while only two strains (EU and E11) carry ISMyma11. Our result also suggested that ISMyma6 and ISMyma7 are the dominant types in the cluster II strains. Remarkably, comparing 1218R with DE4381 (also designated 1218S, which was isolated as a smooth colony variant of 1218R; Small PLC, personal communication), revealed that DE4381 has a higher number of as well as different IS elements than 1218R. Interestingly, both 1218R and 1218S passage strains were likely derived from a common ancestor (TMC1218, Table 1), these results in turn, suggest that a number of IS elements from TMC1218 were lost during passage of 1218R when compared to 1218S.

**Figure 4.**
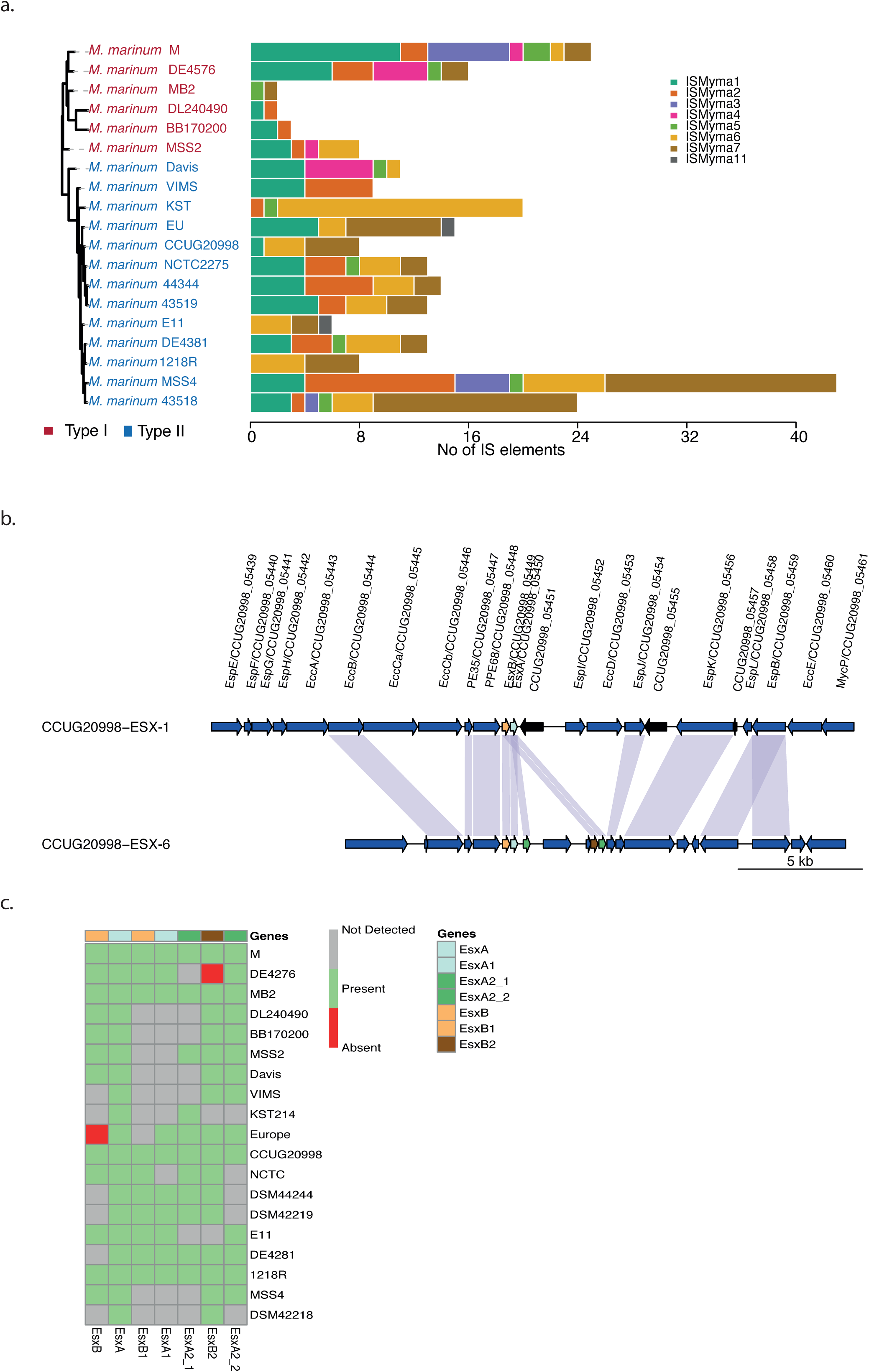
Genomic variations in *Mma* strains. (a) Distribution of IS elements in the *Mma* strains: Bar plot showing copy numbers and distribution of the eight types of IS elements in each of the strains along with their phylogenetic relationship. The phylogenetic tree is the same as shown in Fig 1a. (b) Gene synteny for ESX-1 and the partially duplicated ESX-6 gene clusters. Arrows represent genes and direction of the arrow indicates strand information while vertical connections indicate orthologous genes. Genes are drawn to scale. Color code: blue (ESX-1 related genes) and black (hypothetical protein) arrow mean upstream/downstream of *esxB* and *esxA* gene loci. The regular *esxB* and *esxA* genes are colored as yellow and light green respectively and the connecting light blue shaded vertical line between the arrows indicates homologous genes. (c) Heat map showing presence and absence of esxA and esxB orthologous and paralogous genes in different *Mma* strains.

We also compared the genome-wide distribution of the predicted IS elements in the complete M, CCUG and 1218R genomes. The mapping of predicted IS elements along with whole genome alignment of the three-complete genomes revealed that many divergence regions in the genomes are adjacent to the predicted IS elements (Fig S2).

## Comparative analysis of the gene content: core and auxiliary genes

Overall, the gene content across the different strains is highly conserved with respect to gene synteny and percentage identity. However, several gene clusters present in the M strain are absent in most of the other strains. The numbers of unique genes in the M strain is 277. Of these, 145 (55%) were annotated as hypothetical proteins. These genes are organized in clusters/regions encompassing 9 to 51 genes, and the majority of these are localized in the 3.7 - 4.9 Mb region (Figs 1b and Table S2). Within these regions in the M strain genome, we identified 14 transposase and 19 integrase genes, which raises the possibility that genes within these regions are “readily” mobilized. The unique regions in the M genome also overlap with different phage sequences that add to the diversification of these regions. Furthermore, the presence of prophage sequences raises the possibility that expression of genes within this region(s) is regulated in response to genome rearrangements of prophages, referred to as active lysogeny (Feiner et al. 2015).

Within the unique region near 3.7 Mb, the σ^2997^ gene (MMAR_2997; σ^2997^) was predicted to be present only in the M and NCTC2275 strains [Figs 1 and S3; σ^2997^ is expressed and functional (Pettersson et al., 2015; unpublished)]. Since the σ^2997^ gene is missing in the other *Mma* genomes, it is plausible that it was present in the ancestor but was lost during evolution in the majority of the *Mma* strains.

Many mycobacteria have two potassium (K^+^)-uptake systems, the *trk*- and *kdp*-system (Cholo et al. 2008) and all 19 *Mma* strains were predicted to have genes encoding for the *trk*-system (Table S3). Within one of the unique gene clusters in the M genome we identified the *kdp*-system genes, *kdpABCDEF*. This gene cluster is absent in all the other *Mma* genomes as well as in *M. ulcerans*. This shows that the *kdp*-system is dispensible for growth *in vivo*. It has also been reported that inactivation of the *kdp*-system in *Mtb* increases its virulence (Parish et al. 2003). Whether this also applies to *Mma* warrants further studies.

Core genes constitute the backbone of the genomes, while non-core genes have an impact on the phenotypic variation among different strains. The non-core genes were extracted and plotted in Fig S3b. Of these, 142 genes were predicted to be present in all *Mma* genomes with the exception of BB170200 and DL240490. The predicted number of unique non-core genes varies from 26 (CCUG) to 345 (VIMS) (Fig S3c; Supplementary Table S3). More than 50% of the unique genes in the different genomes were annotated as hypothetical proteins. For many genes, we detected variation in copy numbers comparing the different strains. For example, for genes encoding the dimodular nonribosomal peptide synthase, the tyrosine recombinase (XerD), a few ESX proteins, integrases and transposases. We cannot exclude that some of the genes identified as unique for the 14 draft genomes may be false positives due to the absence of reads in the corresponding loci.

## Duplication of *esxA* and *esxB*

The type VII secretion system was discovered in mycobacteria and ESX-1 genes are major virulence factors for both *Mtb* and *Mma* (Stinear et al. 2008; Houben et al. 2014; Unnikrishnan et al. 2017). As reported for the M strain, all *Mma* strain carry a partial duplication of ESX-1 (the homolog to the prototypical ESX-1 in *Mtb*) gene cluster, resulting in more than one copy of several genes including *esxA*/ ESAT6 and *esxB*/ CFP10 (Figs 4b, c and S4a-d). Interestingly, for the 1218R variant DE4381 (see above), genes positioned upstream of *esxB* in the ESX-1 region have been lost (Fig S4a) but homologues for some of these genes are present in the duplicated ESX-1 region (referred to as ESX-6; Fig S4b; Stinear et al. 2008). Noteworthy, the ESX-1 *esxB* is missing in the Europe and DSM43518 strains, while truncated *esxB* variants are present in MSS2, Davis and DE4576, resulting in shorter protein sequences (Fig S5a). The *esxB* homolog, *esxB*1, in the ESX-6 region in the Europe strain is also missing, while it is present in DE4576 but not in MSS2 and Davis1, which might be due to that they are draft genomes. Hence, for DE4576 the loss of *esxB* could be functionally complemented by *esxB*1 since *esxB* and *esxB*1 are sequentially identical (Fig S5a, b; see also below). Moreover, *esxA* was predicted to be present in all the *Mma* strains (Figs 4b and S4a). For cluster-II members the *esxA* sequence is highly conserved with no sequence variation, while for strains belonging to cluster-I it varies at several positions (Fig S5a). It therefore appears that while the ESX-1 *esxB* gene is dispensable this is not the case for *esxA*.

Comparing the different *Mma* strains the ESX-6 region appears to be more variable than ESX-1 (Figs 4b and S4a, b). In addition to the predicted homologs of *esxB* and *esxA*, *esxB*1 and *esxA*1, we identified the presence of one *esxB* paralog, *esxB*2, and two *esxA* paralogs, *esxA*2 and *esxA*3. The *esxB*2, *esxA*2 and *esxA*3 were predicted to be present in all *Mma* strains with few exceptions, DE4576, KST214 and E11 in the case of *esxB*2 (Fig 4c). EsxB2 is highly conserved and show 54% sequence identity compared to EsxB and EsxB1 (Fig S5). Comparing EsxA and EsxA1 revealed roughly 90% sequence identity, and interestingly, E11 EsxA1 is identical to EsxA1 present in the M strain (Fig S5b). This is in contrast to EsxA (see above) and might possible be due to gene transfer and homologous recombination. On the other hand, the sequences of the EsxA paralogs, EsxA2 and EsxA3, are almost identical across the different strains. However, variation for NCTC and MSS4, where only one paralog was predicted, might be due to that the genomes being draft genomes. As in the case of EsxB2, comparing EsxA2 and EsxA3 with EsxA and EsxA1 revealed lower sequence identities (40-45%) than EsxA and EsxA1 (≈90% sequence identity; Fig S5). Notably, genes within the *Mycobacterium smegmatis* ESX-1 region that influence mating identity have been implicated as having a role for mycobacterial conjugation (Derbyshire and Gray, 2014). However, these genes are not present in the *Mma* or *Mtb* H37Rv ESX-1 regions (Figs 4b and S4a).

Together, these analyses suggest that the duplication of ESX-1 is present in all *Mma* strains and was likely present in the *Mma* ancestor. Furthermore, it appears that the *esxB* and *esxA* genes of the ESX-1 region underwent a second duplication event during evolution to yield an additional ESX region ESX-6 (Figs 4b and S4a; Stinear et al. 2008; Tobias et al. 2013; Kennedy et al. 2014; see discussion).

## Identification of SNVs and mutational hotspots in *M. marinum* strains

Single nucleotide variations (SNVs) were predicted for all the genomes with the M strain as reference using the program MUMmer (Delcher et al. 1999). For cluster-I members, the number of SNVs ranged between 45000 and 56000, while for cluster-II members it is significantly higher, between 70000 and 89000 (Fig 5a). This is consistent with the proposal that the *Mma* strains can be divided into two clusters (see above).

Next, we identified the mutational hotspots in the *Mma* genomes according to Das et al. (2012). Mutational hotspots are genomic regions where the SNV frequencies are much higher relative to the background. One hundred seventy six mutational hotspots were identified in the *Mma* genomes, which corresponds to a frequency of 26.5/Mb (Fig 5b, c). A similar analysis of 20 *Mtb* isolates suggested only 45 mutational hotspots corresponding to 10/Mb (Das et al. 2012; Fig 5c). We therefore determined the hotspot frequencies for three other mycobacteria, *M. avium* subsp. *paratuberculosis* (*MAP*), *M. bovis* (*Mbo*) and *M. phlei* (*Mph*), for which genomic data for several strains are available (see Methods). This analysis revealed that *Mma* carries a higher number of hotspots also compared to these mycobacteria (Figs 5c and S6a-c). Moreover, analysing the two *Mma* clusters separately indicated that the number of mutational hotspots in cluster-I strains is 180, while cluster-II strains have 253 hotspots. However, the average number of SNVs per genomic regions is higher in cluster-I (≈18) than in cluster-II (≈8) consistent with higher divergence among cluster-I members compared to cluster-II members (Fig S7a, b).

**Figure 5.**
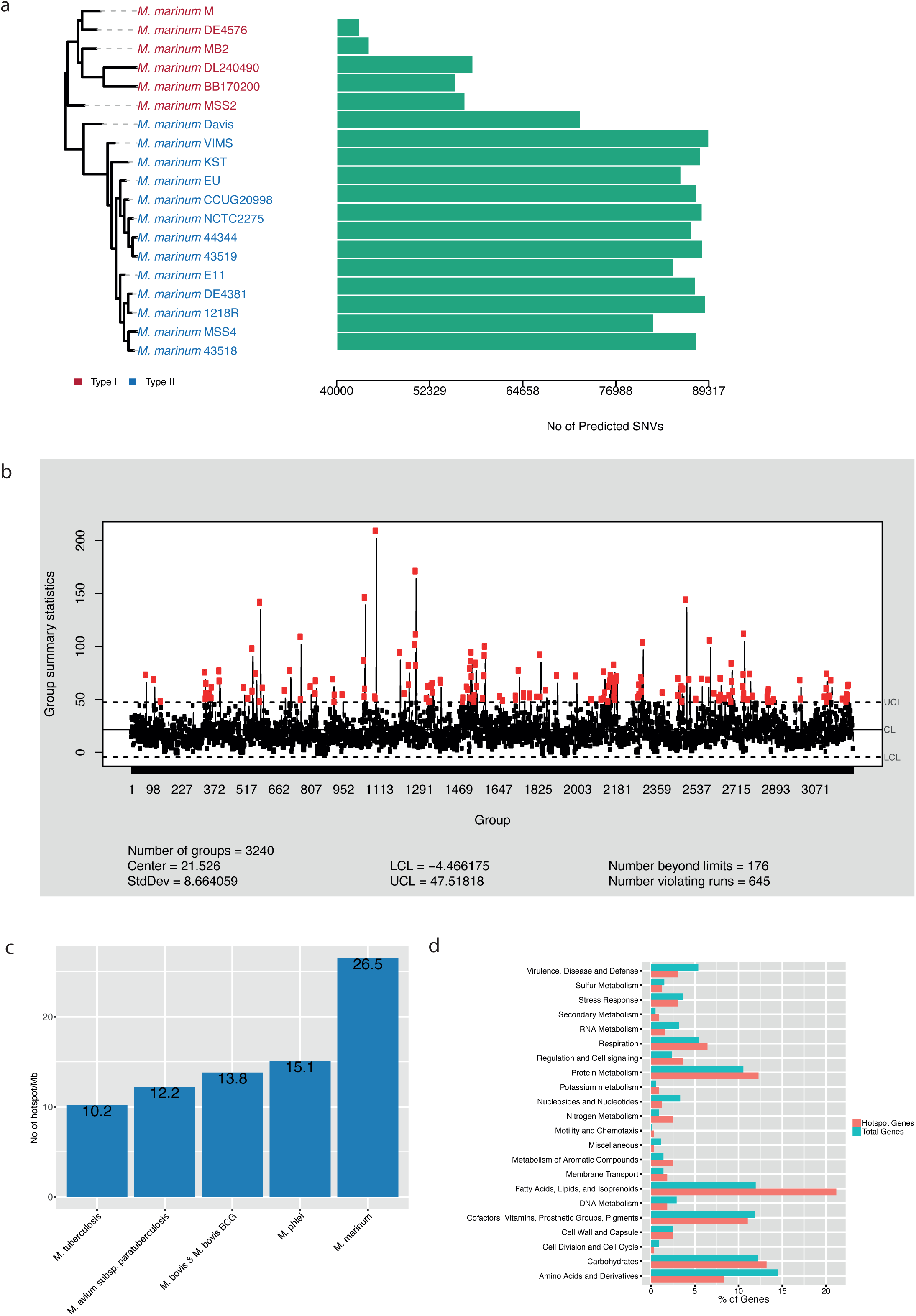
Analysis of mutational hotspots in *Mma*. (a) Bar plot showing the predicted number of SNVs in the different strains compared to the M strain along with their phylogenetic relationship. The phylogenetic tree is the same as in Fig 1a. (b) Shewhart control chart showing the average SNVs frequencies in all the strains. Red and black dots indicate out of control (hotspots) and in-control SNV frequencies, respectively. (c) SNV frequencies per one Mb in different mycobacteria as indicated. (d) Functional classification of genes located in the predicted hotspot regions.

Considering all 19 *Mma* strains, 621 genes map in the hotspot regions. Of these, 300 were annotated as hypothetical genes. The remaining 321 genes were classified into different subsystem categories (Fig 5d), and >20% were predicted to belong to the category "Fatty Acids, Lipids and Isoprenoids". Since *Mma* strains occupy widely different ecological niches this would be consistent with an evolutionary pressure on genes involved in building the outer boundaries.

## Phylogenetic analysis

To understand the phylogenetic relationship between the *Mma* strains, we generated phylogenetic trees based on the 16S rRNA genes using the neighbour-join method with 1000 cycles of bootstrapping. This tree displays three main branches. However, it could not discriminate between closely related strains (Fig 6a). In addition, the 16S rDNA tree is not very robust since many branches have low or zero bootstrap values.

**Figure 6.**
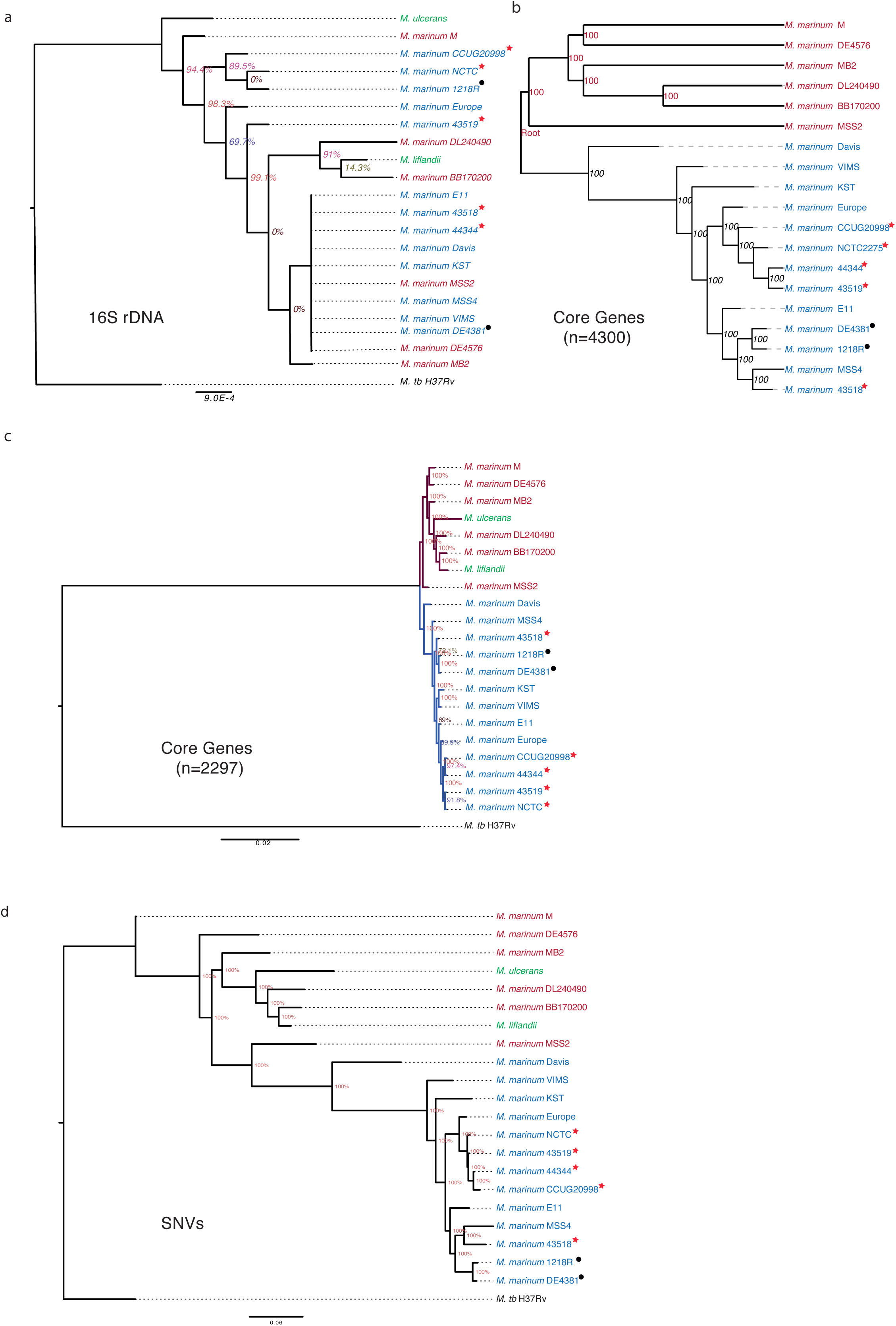
Phylogenetic trees for *Mma*. Phylogenetic trees were based on: (a) 16S rDNA, (b) core genes (n = 4300) present in all *Mma* strains, (c) core genes (n = 2297) in all *Mma* strains, *M. ulcerans* and *M. liflandii* and (d) predicted SNVs compared to the M strain. The percentage values in the nodes represent bootstrap values generated by 1000 cycles. Lab and passage variants derived from the TMC1218 strain (Table 1) are marked with red stars and black circles as indicated.

We previously reported the use of core genes to generate robust phylogenetic trees for other mycobacteria (Das et al., 2015; 2016). Hence, we used the 4300 core genes (see above) present in all 19 *Mma* strains and generated the tree shown in Fig 6b. This tree is supported by high bootstrap values and separates the different strains into two branches/clusters, which is similar to the clustering obtained based on ANI values (cluster-I and -II; cf. Figs 2b and 6b).

*M. ulcerans* and *M. liflandii* are *Mma*’s closest neighbours. Therefore, we were interested in to understand their positions relative to the *Mma* strains. Considering a tree based on 2297 core genes present in all 19 *Mma* strains, *M. ulcerans* and *M. liflandii* cluster together with cluster-I members with MB2, DL240490 and BB170200 as their closest neighbours (Fig 6c). This grouping is in accordance with previous studies that position *M. ulcerans* and *M. liflandii* close to the M strain (Qi et al. 2009; Wang et al. 2015; Doig et al. 2012).

Finally, we generated a tree based on the SNVs identified in the *Mma* strains, *M. ulcerans* and *M. liflandii* (see above and Methods). The resulting tree corroborated the core gene-based phylogeny and clustering of ANI values with few exceptions at the leaf levels (Fig 6d). In agreement with the 2297 core gene tree (Fig 6c) the SNV-based tree suggests that strains belonging to cluster-I are more closely related to *M. ulcerans* and *M. liflandii* than cluster-II members (Fig 6d).

In conclusion, the “M strain lineage” (or M-lineage; cluster-I) members cluster together with *M. ulcerans* and *M. liflandii* and separated from cluster-II (referred to as the “Aronson-lineage”) members. This suggests that members of these two lineages should be classified as two separate mycobacterial species, or at least subspecies. Notably, strains of the “Aronson-lineage” (Table 1, and marked with an * in the figures) that originate from the originally isolated *Mma* strain (Aronson 1926) have diverged, presumably as a result of handling in different laboratories (see discussion).

## Discussion

To trace the evolutionary history and relationships among organisms, the 16S rRNA gene has served as an important biomarker. Specifically it has been useful in providing a reliable phylogenetic tree for bacteria. However, now we have access to large number of bacterial genomes and whole genome comparison can be used to reveal more detailed and better-resolved evolutionary relationships among bacterial species and strains considered phylogenetically unique on the basis of 16S rRNA gene comparison. Consequently, this has expanded the repertoire of genes that can be used to study the evolutionary relationships among bacteria and that have had an impact on their evolution, diversity and use as biotechnological vehicles and documenting the microbial biosphere. Phylogeny based on multiple genes and genomic information have generated more robust trees and in many cases also provided evidence that known species should be considered as separate species or subspecies of known bacteria (Devulder et al. 2005; Das et al. 2015; Loman and Pallen 2015).

We present the genome sequences for 15 *Mma* isolates including the complete genomes of two type strains CCUG20998 and 1218R, both derivatives of the original *Mma* strain isolated by Aronson (1926). Our comparative genomic studies, ANI analysis and phylogenetic trees based on core genes and SNVs, covering 19 *Mma* genomes suggested that the *Mma* strains cluster in two distinct branches, cluster-I and -II. Cluster-I encompasses six strains including the M strain, while the remaining 13 strains constitute cluster-II. In cluster-II we find derivatives of the originally isolated *Mma* strain, *e.g*., the CCUG20998 and 1218R strains. Including *M. ulcerans* and *M. liflandii* revealed that the “M-lineage” (cluster-I) members are their closest neighbours while “Aronson-lineage” (cluster-II) strains are more distantly related. On the basis of these findings, we propose that these two branches should be considered as two species or at least two separate *Mma* subspecies. We suggest that the “Aronson-lineage” should be named *M. marinum* alternatively *M. marinum* subsp. Aronson and the “M-lineage” *Mycobacterium moffett* alternatively *M. marinum* subsp. M (*M. moffett* since it was first isolated at the Moffett Hospital, Universtity of California, San Francisco; Stinear et al. 2008). To distinguish the current “Aronson-” and “M-” strains and for strain identification we further suggest including the name of the strain, *e.g*., *M. marinum* strain 1218R alternatively *M. marinum* subsp. Aronson strain 1218R. Moreover, since available data indicate that *M. ulcerans* evolved from *Mma* (Käser et al. 2007; Doig et al. 2012), our findings indicate that its nearest ancestor belonged to the "M-lineage". In this context, we note that the plasmid fragments present in BB170200 and DL240490, referred to as pMUM003 (Pidot et al. 2008), are highly similar compared to the pMUM001 plasmid present in *M. ulcerans* AY99 (Käser et al. 2007), while plasmids (or plasmid fragments) detected in cluster-II members are different. In addition, plasmid type (I) is only present in strains belonging to the “Aronson-lineage”.

Interestingly, MB2, Europe and DE4576, which all lack plasmids, were isolated as wild outbreak strains in fish; in particular, DE4576 has been shown to be a highly virulent outbreak strain in medaka and zebrafish models (Ennis and Shirreff unpublished), which indicates that plasmids did not play a critical role for virulence for these strains. Moreover, the two *Mma* strains BB170200 and DL240490 were reported to be hyper-virulent due to the production of the mycolactone F toxins *i.e.,* presence of the pMUM003 plasmids (Ucko 2005). Both these mycolactone F producing strains conferred only moderate virulence when compared to the 1218R strain in a controlled infection medaka model (Mosi et al. 2012). In summary, it therefore appears that there is no clear correlation between the presence of plasmid carried in the *Mma* strains studied in this report and virulence in animals; however, 1218R carries a Type I plasmid, while strain DE4381 (a related “smooth” passage variant also called 1218S; see below) has apparently lost this plasmid and may be a product of plasmid segregation. Hence, more detailed genetic and molecular analyses would be required to better document the role that specific plasmids may play in virulence.

The average number of SNVs per region was found to be higher (18.4 vs 7.9; Fig 5c) in members of the “M-lineage” compared to “Aronson-lineage” members. Our data also showed that the “Aronson-lineage” strains CCUG and 1218R (both complete genomes) have two rRNA operons, while the M strain has one (see below; Stinear et al. 2008). Based on that, we predict the presence of two 5S rRNA genes in all the cluster-II draft genomes (one for cluster-I draft genomes) and we assume that all strains in the “Aronson-lineage” carry two rRNA operons. Moreover, as for other bacteria, the *Mma* pan-genome is open (*B_pan_* = 0.47; Fig 3a; see also *e.g.,* Lefébure and Stanhope 2007; Lukjancenko et al. 2010; Jacobsen et al. 2011; Gordienko et al. 2013; Vernikos et al. 2015). Its size amounts to 8725 genes. Of these, roughly 50% constitute the core genome. However, comparing the pan-and core-genomes of the “M-” and “Aronson-” lineages revealed that the pan-genome for the “M-lineage” is larger, while its core-genome is smaller (Fig 3c). It should be noted that the pan- and core-genome sizes vary for individual species within the *Streptococcus* genus (Lefébure and Stanhope 2007). Taken together, these findings suggest that the “M-lineage” members show higher diversity compared to members belonging to the “Aronson-lineage”. As such, support the notion that the two lineages should be considered as separate species (or subspecies).

It has previously been reported that mutational hotspots in *Mtb*, expressed as SNVs per genome size (Mb), cluster in certain genomic regions (Das et al. 2012). We provide data that the number of mutational hotspots is significantly higher for the 19 *Mma* strains compared to other mycobacteria (*Mtb*, *MAP*, *Mbo* and *Mph*; Fig 4c). A major fraction of these hotspots were mapped to genes involved in fatty acid, lipid and isoprenoid metabolism. Many of these genes play important roles in building and altering the outer boundaries in response to environmental changes, consistent with that *Mma* inhabits different ecological niches. In this context, we note that two strains (MSS2 and MSS4) belonging to different lineages, MMS2 was isolated from an infected fish cultivated from a striped bass aquaculture facility and MMS4 from a patient working at the same fish farm (Ostland et al. 2008). This suggests that the two strains occupy the same ecological niche. One expectation would be that the same strain causing disease in the fish would also be the one infecting the human. However, our genomic analysis showed that this was not the case. Rather, this might be related to different strains having a selective advantage for infecting different hosts. Alternatively, this finding could be random and insignificant.

The sole rRNA operon (*rrnA*) in the M strain is located downstream of *murA* and upstream of the *apt* gene (coding for O_6_-alkylguanine DNA alkyltransferase). In general, rapidly growing mycobacteria harbour two rRNA operons, *rrnA* and *rrnB*, and these are located downstream of the *murA* and *tyrS* genes, respectively (Menendez et al. 2002). However, *rrnA* and *rrnB* in the cluster-II members are located next to each other separated by a duplicated copy of the *apt* gene (Fig 1b). This suggests that the presence of two *rrn* genes in cluster-II members likely is the result of a duplication event. Since the closest neighbours of *Mma*, *i.e*. *Mtb*, *Mycobacterium kansasii* and *Mycobacterium gastri*, are equipped with one rRNA operon it is likely that the duplication occurred after the “M-” and “Aronson-” lineages diverged. However, we cannot exclude that their ancestor had two rRNA operons and that one was lost after the two lineages diverged. In this context, we note that the M and CCUG strains grow with similar rates in 7H10 media (Ramesh, unpublished) consistent with that the number of rRNA operons does not affect the growth rate (see *e.g.* Gonzalez-Y-Merchand et al. 1998; 1999; Menédez et al. 2005).

IS elements have a key role in generating diversity among bacteria. As such, IS elements can be used as biomarkers for strain identification, epidemiological tracking and predicting spread of antibiotic resistance (Eisenach 1994; Gunisha et al. 2001; Warren et al. 2009). We identified the presence of eight known IS elements (ISMyma1-7, 11) in all 19 *Mma* strains. Their distribution and copy number varied among the different strains with MSS4 having the highest number, 44 IS elements (Fig 4). Hence, these data open up for the development of strain specific *Mma* probes for use in clinical settings that relate to, *e.g.,* fisheries and aquariums. For example, using probes to determine whether an infection is caused by MSS2 or MSS4 (see above). In this context, we note that more than four ISMyma3 copies are present in the M and MSS4 strains, which were both originally isolated from human patients.

Of specific interest is the ESX-1 region and in this context the variants, 1218S and 1218R, which are passage variants derived from the TMC1218 lineage (Table 1). We have studied the properties of the DE4381 (”1218S”) smooth colony variant, and our data showed that it only conferred a modest (≈ten-fold) reduction of virulence when compared to the rough colony variant 1218R using the established Japanese medaka infection model system (Broussard and Ennis, 2007; Ennis and Cheramie, unpublished). The DE4381 strain carries a deletion encompassing ten genes within the ESX-1 interval, including the loss of *esxB* and nine other well-defined ESX-1 genes (Fig S4a). The loss of one of these genes, *eccA*1 (which corresponds to Rv3868 in *Mtb*H37Rv), has been reported to confer large pleiotropic effects on virulence (Gao et al. 2004). In keeping with this, a *Mma* Δ*eccA*1 mutant strain displays a substantial reduction (>1000-fold) in spread and colonization upon infection of Japanese medaka (Mallick and Ennis, unpublished). We are therefore endeavouring to better characterize these unexpected subtle effects on virulence that was conferred by this ten-gene deletion and this may suggest that the DE4381 (1218S) strain carry suppressor mutations for virulence. We expect that by comparing other smaller rearrangements and mutational changes between the 1218R and DE4381 (1218S) strains, and differences in transcriptional profiles, will gain insights into the compensational mutations that presumably result in this partial suppression of virulence associated with the DE4381 strain. In this context we also note that absence of ESX-1 genes have been discussed to be associated with changes in colony morphologies such as exhibition of smooth or rough colony morphology (Bosserman and Champion 2017).

To conclude, our study provides insight into the diversity of *Mma* strains. Several factors contribute to this diversity, such as the presence of phage and IS elements as well as plasmids. The diversity is also expressed in terms of higher numbers of mutational hot spots compared to other mycobacteria. Together this emphasises that the *M. ulcerans*-*M. marinum* complex, MuMC, constitute a group of bacteria to identify factors and study their importance for bacterial evolution (see Röltgen et al. 2012).

## Methods

### Strain information and cultivation

Information about the *Mma* strains is compiled in Table 1. DSM44344, DSM43518 and DSM43519 were purchased from DSMZ (Deutsche Sammlung von Mikroorganismen und Zellkulturen GmbH). MSS2 was isolated from Hybrid striped bass during an outbreak in Mississippi, USA in an aquaculture facility, and MSS4 from a lesion of a human patient who worked in this facility. VIMS9 was isolated from wild striped bass in Virginia, USA and Davis from farmed striped bass in Davis, California, USA. The different strains were cultivated as described elsewhere (Das et al. 2015; 2016).

### DNA sequencing and assembly

Complete genomes of the *Mma* type strains, CCUG20998 and 1218R were sequenced using PacBio technology. The remaining 13 strains were sequenced on HiSeq2000 (Illumina platforms) at the SNP@SEQ Technology Platform, Uppsala University. Genomic DNA was isolated and prepared for sequencing as described elsewhere (Das et al. 2015; 2016).

PacBio sequencing reads with an average length more than 10,747 bp and read depth of around 100x were assembled using the SMRT-analysis HGAP3 assembly pipeline (Chin et al. 2013), polished using Quiver (Pacific Biosciences, Menlo Park, CA, USA) and generated single scaffolds for CCUG20998 and 1218R.

For the Illumina sequencing reads, a total of 12 million short reads was generated for each strain with an average read length of 100 nucleotides (Table 1). Filtering of the short reads was done to remove low quality reads and ambiguous bases. *De novo* assembly of the short reads was done using the A5 assembly pipeline (version 1.05; Tritt et al. 2012). Final genomes consist of contigs of more than 200 bases. Scaffolds were re-ordered using MAUVE with the CCUG20998 genome as reference.

### Genome Annotations

All the genomes, including the available M, Europe, E11 and MB2 genomes, were annotated using the Prokka pipeline (Seemann 2014), which uses Prodigal (Hyatt et al. 2010) to predict coding sequences (CDS). tRNA and rRNA genes were predicted by Aragorn (Laslett and Canback 2004) and RNAmmer (Lagesen et al. 2007). Annotated genes were functionally classified using the RAST Subsystem (Aziz et al. 2008).

### Identification of plasmid fragments and foreign DNA

Plasmid fragments were identified by pairwise alignments of the scaffolds with sequences from the NCBI plasmid database (ftp://ftp.ncbi.nlm.nih.gov/genomes/Plasmids/).

Prophage sequences were identified using concatenated ordered scaffolds for all the genomes in the PHAST server (Zhou et al. 2011).

### Prediction of IS elements

IS elements were predicted for the complete genomes using the ISsaga server (Varani et al. 2011). All the predicted IS elements were used as reference to identify IS elements in the draft genomes using ISmapper (Hawkey et al. 2015). ISmapper uses raw reads and reference IS elements and predicts possible positions of the reference IS elements in the genome enquired.

### Identification of orthologous genes

Homologous coding sequences were identified using an all-versus-all BLAST search of the protein sequences from all 19 strains. Orthologous genes were predicted using PanOCT (v3.23) from the BLAST output. PanOCT follows two criteria to consider a gene as orthologous, sequence homology and gene synteny. All necessary PanOCT input files were generated using in-house shell scripts, and PanOCT was executed to detect the genomic differences based on coding regions.

### Identification of SNVs and mutational hotspots

Whole genome alignments were performed in a pairwise manner using MUMmer (Delcher et al. 1999). SNVs were identified using the "show-snps" program of the MUMmer package. Single nucleotide insertions/deletions were filtered out, and only SNVs were used for further analysis. Mutational hotspots were identified using Shewhart Control Chart, as described by Das et al. (2012). Briefly, the genome of the M strain was divided into non-overlapping windows of 2000 bases and the average number of SNVs in each of the windows was determined. The average SNV values were subsequently used in Shewhart Control Chart for the prediction of hotspots. Mutational hotspots were identified for *Mma* (n = 19), *Mbo* (n = 28), *Mph* (n = 5), and *MAP* (n = 23). Mutational hotspots for cluster-I and cluster-II were identified separately using the genome sequences of the M and CCUG strains as references for cluster-I and cluster-II, respectively, and follow the procedure as described above.

## Data deposition

This Whole Genome Shotgun project has been deposited at DDBJ/ENA/GenBank under the project PRJNA414948, PRJ414525.

## Acknowledgements

We thank our colleagues for discussions. Sequencing was performed by the SNP@SEQ Technology Platform in Uppsala, which is part of the Science for Life Laboratory at Uppsala University and supported as a national infrastructure by the Swedish Research Council. The computations were performed on resources provided by SNIC through Uppsala Multidisciplinary Center for Advanced Computational Science (UPPMAX) under Project b2011072. This work was funded by the Swedish Research Council (M and N/T), the Swedish Research Council for Environment, Agricultural Sciences, and Spatial Planning (FORMAS), and Uppsala RNA Research Center (Swedish Research Council Linneus support).

## Author contributions

LAK and DGE conceived the study. S Das designed and performed the bioinformatics computations and PRKB bioinformatics analysis. S Das, DGE and LAK analyzed and interpreted the data. AM, MC, LS and TD maintained, cultivated and prepared DNA from different *Mma* strains. BMFP and DGE generated culture extracts and DNA isolation. S Das, S Dasgupta, MC, DGE and LAK wrote the manuscript. All authors read and approved the final version of the manuscript.

## Table Legends

**Supplementary Table S1** Summary of the genomic features and annotation for *Mma* strains used in this study.

**Supplementary Table S2** Detailed information about predicted phages in the different *Mma* genomes.

**Supplementary Table S3** List of common and unique genes and their functions in the various *Mma* strains.

## Figure Legends

**Figure S1 Genome sequencing reads and assembly statistics** (a) Bar plots showing number of raw reads, coverage, N50 and number of scaffolds in the 14 *Mma* genomes sequenced using Illumina sequencing platform.

(b) Plots showing average read lengths, coverage, and raw reads for the 1218R and CCUG20998 genomes sequenced using Pacific Biosciences technology.

**Figure S2 Genome wide distribution of IS elements in complete genomes** Circos plot of the genome alignment of three complete genomes: M, 1218R and CCUG20998 represented by blue, green and orange arcs, respectively. Lines connecting the arcs show homologous regions in the genome while white gaps between the lines indicate unique genomic regions. Predicted IS elements are marked with black bars in the corresponding genomes. Scales outside the arcs shows the lengths of the genomes.

**Figure S3 *Mma* M strain orthologous and non-core genes** (a) Circos plot showing the presence of protein coding genes in 18 *Mma* strains compared to the M strain. The outer track represents the genome for the M strain with a size scale. Radial black lines with blue fill mark genes in the M strain. Each circular track represents one genome and the number corresponds to the strain name in the legend. Coloured radial blocks represent orthologous genes in the corresponding genome and colour intensity indicates percentage identity at the protein levels. The white blocks indicate that no orthologs were identified. (b) Heat map showing presence (red) and absence (white) of orthologous genes in the different *Mma* strains. Clustering of the values was done using hierarchical clustering. (c) Stacked bar plot showing the percentage of unique and non-unique genes in all *Mma* strains.

**Figure S4 Gene synteny plot of ESX-1 regular and partially duplicated gene clusters in different *Mma* strains** Gene synteny plot showing (a) complete ESX-1 and (b) partially duplicated ESX-6 gene clusters in all the *Mma* strains. Arrows represent genes and their direction indicates strand information while vertical connections indicate orthologous genes. Genes are drawn to scale. Color code: blue (ESX-1 related genes) and black (hypothetical protein) arrow mean upstream/downstream of *esxB* and *esxA* loci. The regular *esxB* and *esxA* genes are coloured as yellow and light green respectively and the connecting light blue shaded vertical line between the arrows indicate homologous genes. The *Mtb*H37Rv genome was re-annotated (marked as Rv_Prokka) using the same approach as used for annotation of the other genomes. For comparison we included the annotation of *Mtb*H37Rv (marked Mtb in the figure) used by Stinear et al. (2008). Comparing the two annotations revealed that the original EspI/RVBD_3876 gene is annotated as two genes as predicted to be the case in the *Mma* genomes. In (b) the regular *esxB* and *esxA* orthologs are coloured in yellow and light green while the respective paralogs are coloured in brown and green. The genes marked in shaded light blue indicate homologous genes. (c) Heat map showing presence and absence of esxA and esxB orthologous and paralogous in 1218R. (d) Similar to (c) but for the M strain.

**Figure S5 Multiple sequence alignments: EsxA and EsxB and their orthologs in the *Mma* strains** (a) Top, alignment showing EsxA and its orthologous genes while the bottom alignment represents EsxB and its orthologs in the Mma strains.

(b) Similar to a, top alignment for EsxA1 and bottom showing EsxA2.

(c) Similar to a, b and the top alignment show EsxB1 and bottom EsxB2.

**Figure S6 Analysis of mutational hotspots in other mycobacteria** Shewhart control chart showing average SNVs frequencies for different mycobacteria as indicated. Red and black dots mark out of control (hotspots) and in-control SNV frequencies, respectively.

(a) 26 strains of *Mbo* (covering both *M. bovis* and *M. bovis* BCG genomes).

(b) Five *Mph* strains (Das et al., 2016).

(c) 23 MAP strains

The different genomes were extracted from the NCBI database.

**Figure S7 Analysis of mutational hotspots in members of cluster-I and cluster-II** Shewhart control charts showing the average SNVs frequencies for: (a) cluster-I strains and (b) cluster-II strains. Red and black dots indicate out of control (hotspots) and in-control SNV frequencies, respectively.

